# OT-knn: a neighborhood-aware optimal transport framework for aligning spatial transcriptomics data

**DOI:** 10.64898/2026.02.19.706743

**Authors:** Jia Song, Qunhua Li

## Abstract

Spatial transcriptomics (ST) measures gene expression while preserving spatial context within tissues, enabling detailed characterization of tissue organization. As ST technologies advance, aligning datasets across tissue sections, individuals, platforms, and developmental stages has become increasingly important but remains challenging due to sparse expression, biological heterogeneity, and geometric distortions between slices.

We introduce OT-knn, a method for ST alignment that integrates local neighborhood information within an optimal transport framework. Rather than relying solely on single-spot expression, OT-knn reconstructs each spot using its spatial *k*-nearest neighbors, capturing microenvironment context that is more robust to noise and variability. These representations are then used to derive probabilistic correspondences between slices. We evaluate OT-knn using simulated data with known ground-truth alignment and real datasets from multiple ST platforms, including human dorsolateral prefrontal cortex data (10x Genomics Visium), mouse brain aging data with both within-donor and cross-donor comparisons (MERFISH), a multi-stage axolotl brain dataset (Stereo-seq), and a cross-platform analysis using mouse embryo datasets profiled by Stereo-seq and seqFISH. Across these settings, OT-knn achieves accurate and robust alignment, particularly in the presence of spatial deformation, donor heterogeneity, developmental variation and technological differences.

**Author summary:** Understanding how cells are arranged within tissues is important for studying how organs develop, age, and respond to disease. Spatial transcriptomics is a new technology that measures gene activity while keeping track of where each measurement comes from in the tissue. However, comparing data from different tissue slices, different individuals, or different time points is difficult. Tissues can change shape during preparation, gene measurements can be noisy, and natural biological differences exist across samples.

In this work, we develop a method to better match corresponding regions across tissue slices. Instead of looking at each location by itself, we also consider information from its surrounding neighborhood. This provides a more stable and informative view of the tissue and makes it easier to match similar regions, even when the data are imperfect or the tissue shapes differ. We test our method on both simulated and real datasets from human, mouse, and axolotl brain tissues. Our approach enables more reliable comparisons of spatial gene activity across samples, which can help researchers study development, aging, and disease.

## Introduction

Spatial transcriptomics (ST) enables the measurement of gene expression while preserving the spatial organization of cells within tissues [1]. By integrating transcriptomic profiles with spatial context, ST provides critical insights into cellular composition, tissue architecture, and cell–cell interactions that are inaccessible to dissociated single-cell assays. As a result, ST technologies have been widely adopte across diverse biomedical fields, including neuroscience [2–7], cancer [1,8–11] immunology [12–14], developmental biology [7, 15–17], and reproductive biology [18].

Over the past decade, a wide range of ST technologies have emerged, broadly categorized into sequencing-based and imaging-based approaches. Sequencing-based platforms, such as Slide-seq, Visium, Visium HD, and Stereo-seq [15, 19–22], use spatial barcoding and next-generation sequencing to achieve genome-wide transcriptomic coverage, typically at moderate spatial resolution. In contrast, imaging-based methods, including MERFISH, seqFISH, and Xenium [23–25], directly image RNA molecules to achieve much higher spatial resolution but measure fewer genes and require more specialized instrumentation. These differences in spatial resolution, gene coverage, and data characteristics across platforms make alignment challenging and highlight the need for methods that are sufficiently general to perform reliably across diverse ST technologies.

A central challenge in spatial transcriptomics analysis is the alignment of ST datasets across tissue sections, samples, experimental conditions, technological differences, or developmental stages. Accurate alignment enables downstream tasks such as comparative analysis, cross-sample integration, and three-dimensional reconstruction of tissue architecture. However, alignment remains difficult due to several factors. First, ST data are inherently sparse and high-dimensional, with substantial dropout that weakens transcriptomic similarity between corresponding spots [26]. Second, biological variation across samples—arising from inter-individual differences, changes in cell-type composition, or developmental progression—can obscure true anatomical correspondences [26, 27]. Third, technical and experimental factors introduce additional variability, including batch effects, platform-specific biases, and geometric distortions caused by tissue handling, sectioning, or deformation. Even adjacent tissue slices may exhibit nonlinear spatial warping, expansion, or shrinkage, violating assumptions of rigid geometry and further complicating alignment [26, 28].

Several computational methods have been developed to address ST alignment. PASTE [29] uses a fused Gromov–Wasserstein optimal transport framework to jointly model transcriptional and spatial similarity. DeST-OT [30] extends this approach by employing semi-relaxed optimal transport to better accommodate developmental changes. STalign [31] applies large deformation diffeomorphic metric mapping to register spatial gene expression patterns, while GPSA [32] uses Gaussian processes to embed samples into a shared latent coordinate system. More recently, SLAT [33] combines graph neural networks and adversarial matching to improve correspondence across slices. Spateo [34] uses probabilistic point-set registration that integrates molecular profiles and spatial coordinates to support partial, non-rigid, and multi-slice alignment. Despite these advances, robust alignment across highly heterogeneous, noisy datasets, particularly when slices differ substantially in geometry or developmental state, remains an open problem.

In this work, we introduce OT-knn, a spatial transcriptomics alignment method that integrates local neighborhood information within an optimal transport framework. The key motivation behind OT-knn is that the local neighborhood of a spot provides a more stable and biologically meaningful representation than single-spot expression alone. While individual spots are often noisy or incomplete, their surrounding spatial context captures local tissue organization and cell–cell relationships that are more robust to technical noise and biological variability. OT-knn leverages this idea by reconstructing each spot’s expression profile from its *k* nearest neighbors, embedding local spatial structure before performing probabilistic alignment across slices using optimal transport.

We evaluate OT-knn on both simulated and real datasets spanning multiple platforms and biological settings. Simulated data are generated from a human intestine spatial transcriptomics dataset [35] to enable controlled assessment under spatial distortion and expression perturbation. Real-data evaluations are performed on a human dorsolateral prefrontal cortex (DLPFC) dataset [5] from 10x Genomics Visium [21], a within and cross-donor mouse brain dataset [7] generated by MERFISH [23] that allows evaluation of both within-donor and cross-donor alignment, a multi-stage axolotl brain dataset [17] generated using Stereo-seq [15], and a cross-platform mouse embryo datasets profiled by Stereo-seq [15] and seqFISH [24, 36]. Across these diverse scenarios, OT-knn consistently achieves accurate and robust alignment, particularly in settings involving geometric distortion, expression variability, donor heterogeneity, technological differences, and developmental dynamics.

## Results

### Overview of OT-knn

OT-knn is a computational method for aligning spatial transcriptomics (ST) data across tissue slices by jointly leveraging gene expression profiles and spatial environment. It represents each spot as a weighted aggregation of its own and its *k*-nearest neighbors’ gene expression, thereby embedding the spot within its local tissue environment. These representations are then aligned using an optimal transport (OT) framework, which produces a probabilistic mapping that quantifies correspondences between spots. This mapping supports both soft alignments, which is useful for uncertainty modeling, and one-to-one matching when strict correspondence is required.

Specifically, OT-knn takes as input the gene expression profiles and spatial coordinates from two slices (Fig. 1). It first selects the top 5,000 highly variable genes shared between slices (see Methods). For each spot, OT-knn reconstructs its expression by computing a distance-weighted average of its *k* nearest neighbors, giving higher weights to closer neighbors. It then solves a balanced OT problem to minimize transcriptional dissimilarity between the reconstructed profiles, yielding a probabilistic alignment matrix. One-to-one correspondences can be obtained by selecting the spot pairs with the highest transport probabilities.

**Fig 1.**
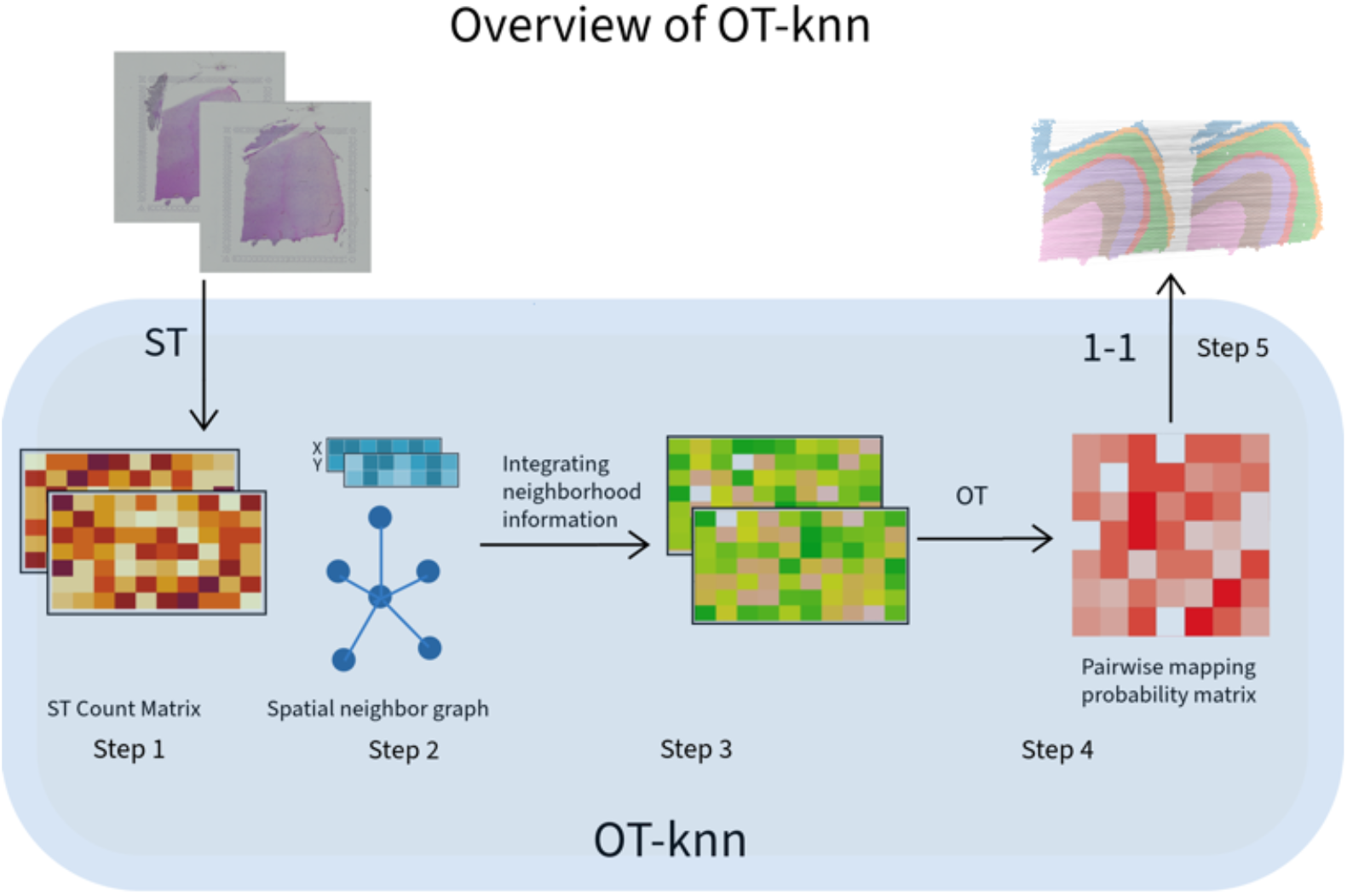
Overview of OT-knn. Given the gene expression and spatial coordinates of ST data from two tissue slices, OT-knn (1) selects shared highly variable genes, (2) builds spatial kNN graphs, (3) refines each spot’s gene expression profile by a distance-weighted average of its *k* neighbors, (4) computes a probabilistic alignment with optimal transport, and (5) extracts best-matching pairs.

For analyses involving multiple slices, OT-knn can be applied sequentially to adjacent slice pairs, and the resulting correspondences can be linked across slices to support multi-slice integration, domain identification, and three-dimensional reconstruction.

### Simulation-based evaluation under spatial and transcriptional perturbations

To understand how alignment methods respond to controlled perturbations across slices, we first evaluated them using simulated spatial transcriptomics (ST) data with known spot-to-spot correspondence. This framework enables direct quantification of alignment accuracy and examination of method-specific error patterns, which is not possible for real tissue sections with unknown ground-truth correspondence.

We used an adult human intestine ST slice profiled with the 10x Genomics Visium platform [21, 35] as the reference and generated a second, perturbed slice under four scenarios (see Methods). The first three controlled scenarios isolated the effects of (1) nonlinear spatial distortion, (2) gene expression variation, and (3) their combination. The fourth, more challenging scenario combined global slice misalignment, local non-rigid deformation, gene expression variation, and spot dropout.

We compared OT-knn with PASTE [29], DeST-OT [30], SLAT [33], and Spateo [34]. Performance was assessed using best-matching accuracy and mapping probability accuracy (Fig. 2b; see Methods), and one-to-one matching patterns were visualized to examine the spatial distribution of alignment errors (Fig. 2c).

**Fig 2.**
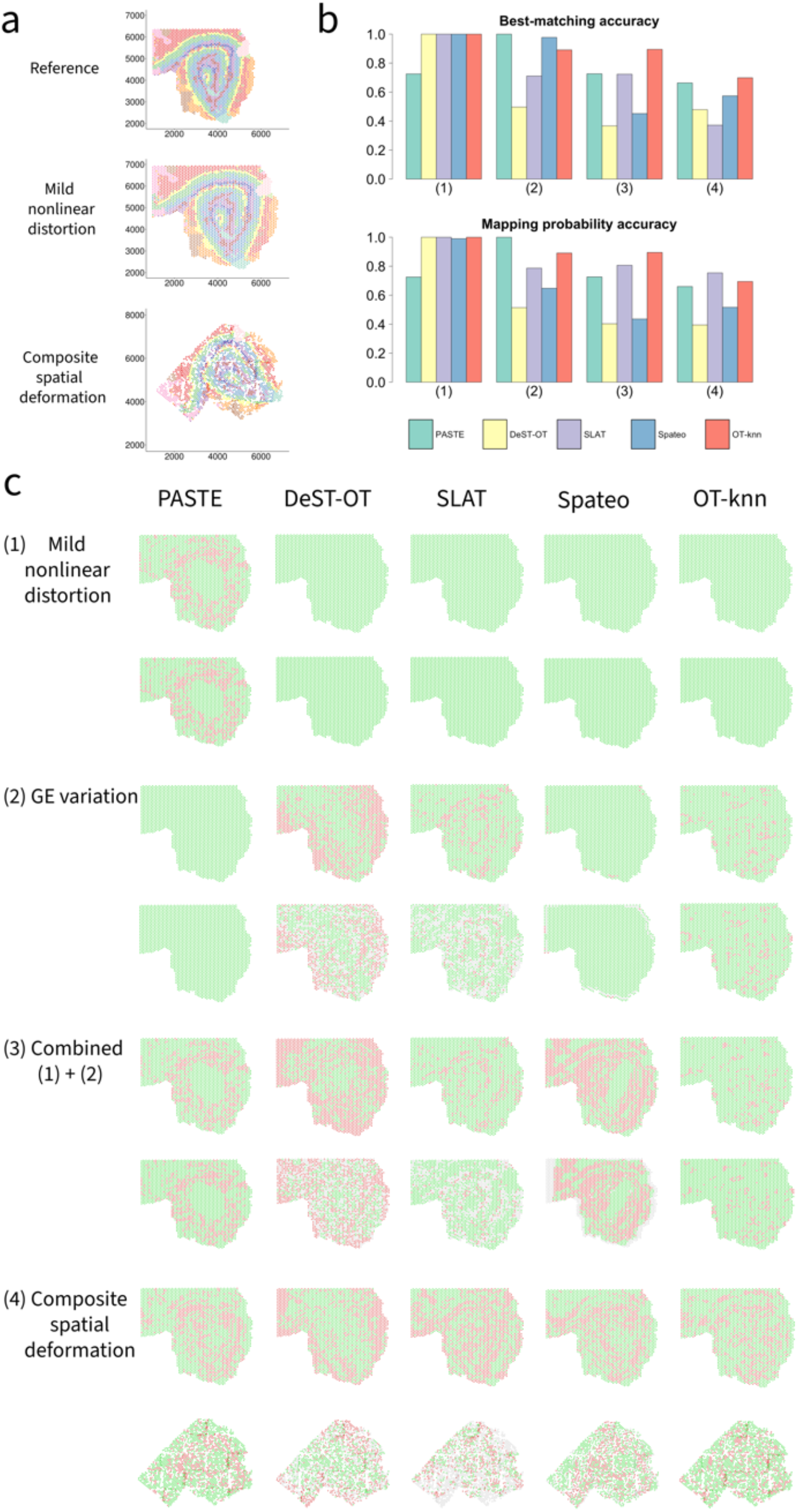
Performance comparison under controlled perturbations. **a**. Reference slice (top), mild nonlinear distortion slice (middle), and composite spatial deformation slice (bottom), colored by clusters obtained using Seurat. **b**. Best-matching accuracy (top) and mapping probability accuracy (bottom) under four simulation scenarios (1-4) for PASTE, DeST-OT, SLAT, Spateo, and OT-knn. **c**. One-to-one matching results. Correctly aligned spots are shown in green, misaligned spots in red, and unaligned spots in gray. For each scenario, the top row shows the reference slice and the bottom row shows the simulated slice.

Under nonlinear spatial distortion, DeST-OT, SLAT, Spateo, and OT-knn maintained high alignment accuracy, whereas PASTE exhibited substantial misalignment, consistent with its stronger reliance on spatial coordinates. Under gene expression variation, PASTE and Spateo remained accurate, whereas DeST-OT and SLAT showed reduced accuracy and left many spots unmatched in one slice. OT-knn exhibited a modest decrease in performance but consistently outperformed both methods and aligned all spots. Under combined perturbation, DeST-OT and Spateo degraded further, indicating only partial robustness to spatial deformation and expression noise, while OT-knn achieved the highest overall accuracy.

These results reveal distinct sensitivities among the evaluated methods. PASTE’s fused Gromov–Wasserstein formulation encourages preservation of the within-slice spatial distance structure; nonlinear deformation alters this structure and can therefore reduce alignment accuracy. DeST-OT was relatively tolerant of spatial distortion alone but was more strongly affected when gene expression variation was introduced, particularly under the combined perturbation. (Fig. 2c). SLAT left many spots unmatched in one slice even when the two slices had equal numbers of spots and identical spatial geometry. This finding suggests that differences in slice size are not the sole source of the asymmetric matching reported for SLAT [33] and is consistent with its directional matching procedure, which does not impose balanced mass-conservation constraints. Spateo achieved high best-matching accuracy when spatial distortion or gene expression variation was introduced separately. However, its mapping probability accuracy decreased in the presence of gene expression variation, and its overall performance declined further when spatial and transcriptional perturbations were combined.

The first three scenarios isolate method sensitivity to spatial and transcriptional variation. We next evaluated performance under the fourth, composite scenario, in which multiple sources of variation were introduced simultaneously. Under the composite perturbation, OT-knn achieved the highest best-matching accuracy among all evaluated methods (Fig. 2b). The one-to-one matching results also showed fewer mismatched and unmatched spots for OT-knn than for the competing methods (Fig. 2c), demonstrating robust performance when global misalignment, local deformation, transcriptional variation, and spot dropout occurred simultaneously.

Overall, these results suggest that incorporating local spatial context into spot-level expression representations improves robustness to spatial deformation, transcriptional variation, and spot dropout.

### Human DLPFC data (10X Genomics Visium)

We applied OT-knn to human dorsolateral prefrontal cortex (DLPFC) spatial transcriptomics data [5] generated using the 10X Genomics Visium technology [21]. The dataset contains 12 tissue slices from three donors, with each slice manually annotated into six neocortical layers and white matter (WM). Each donor, denoted Sample I, II, or III, contains four anatomically ordered slices (A–D). The A–B and C–D pairs are separated by 10 *µ*m, whereas the B–C pair is separated by 300 *µ*m (Fig. 3a).

**Fig 3.**
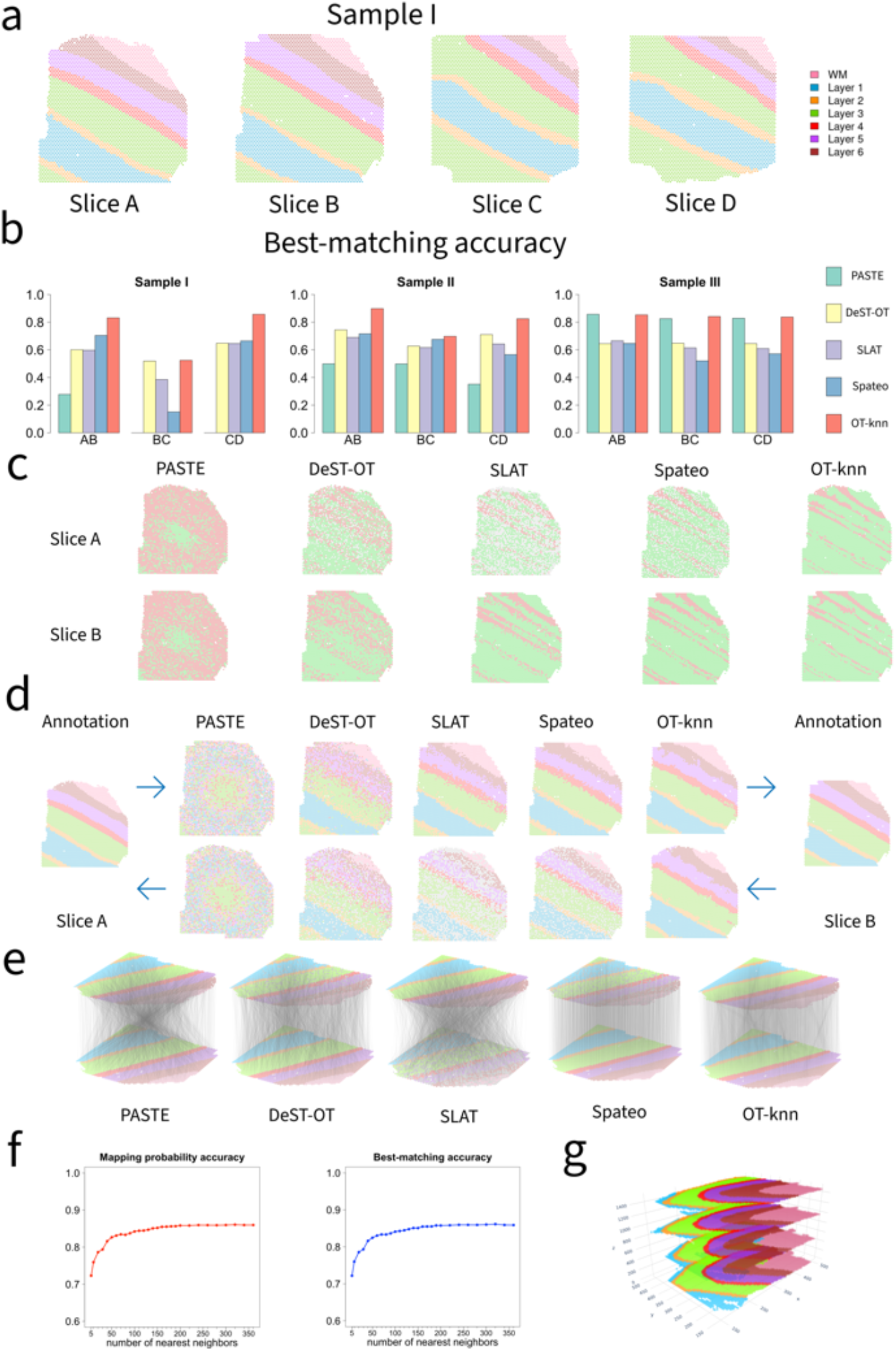
Performance comparison for human DLPFC dataset. **a**. DLPFC Sample I’s four tissue slices, colored according to the manually annotated layers by Maynard et al. [5]. **b**. Best-matching accuracy of consecutive slices across the three samples for PASTE, DeST-OT, SLAT, Spateo, and OT-knn. **c**. Best-matching results for the first pair of Sample I. Correctly aligned spots are shown in green, misaligned spots in red, and unaligned spots in gray. **d**. Domain inference results for the first pair of Sample I by PASTE, DeST-OT, SLAT, Spateo, and OT-knn. **e**. Visualization of the alignment for the first slice pair of Sample I. Spots are colored according to their annotated cortical layers, and unaligned spots are shown in gray. Connection lines represent the best-matching spots across the two slices. **f**. Mapping probability accuracy and best-matching accuracy across different numbers of nearest neighbors for the middle pair of Sample II. **g**. Stacked 3D alignment of the four tissue slices of Sample III after alignment with OT-knn.

#### Pairwise alignment performance

We first evaluated pairwise alignment between each pair of consecutive slices (A–B, B–C, and C–D) within each donor, yielding nine alignments in total. OT-knn achieved the highest best-matching accuracy in eight of the nine comparisons and was a close second in the remaining comparison (Fig. 3b). For the closely spaced A–B and C–D pairs in Samples I and II, as well as all three pairs in Sample III, OT-knn achieved accuracies ranging from 82.5% to 89.8%. Its performance decreased for the more widely separated B–C pairs in Samples I and II, to 52.4% and 69.8%, respectively. A similar decrease was observed for all methods, consistent with greater anatomical differences between these slices.

PASTE performed comparably to OT-knn in Sample III (82.5-85.7%) but was substantially less consistent in Samples I and II (0-49.9%). DeST-OT achieved accuracy similar to OT-knn for the B–C pair in Sample I (51.9%) but was lower for most other comparisons (60-74.5%). SLAT and Spateo generally showed lower best-matching accuracy across the nine alignments (38.5-69% and 15.1-71.6%, respectively).

Mapping probability accuracy showed a similar overall pattern (S2 Fig). For the closely spaced pairs, OT-knn consistently exceeded 82.5% (82.6–90%), compared with 74.7-89.2% for Spateo, 74.2–87% for SLAT and 66.5–83% for DeST-OT. PASTE again performed comparably in Sample III (82.6-85.7%) but substantially worse in Samples I and II (24.7-49.5%). For the more widely separated B–C pairs in Samples I and II,

OT-knn, DeST-OT, and SLAT maintained moderate and relatively consistent accuracy (52.5–72.1%), whereas PASTE was substantially lower (20.8% and 49.9%) and Spateo showed highly variable performance across the two samples. In Sample III, OT-knn and PASTE consistently achieved the highest mapping probability accuracy across all three slice pairs, remaining above 82.6%, whereas DeST-OT, SLAT, and Spateo generally showed lower and more variable performance (72.6-79.1%).

#### Alignment patterns and label transfer

Visualization of one-to-one matches for the A–B pair of Sample I showed that alignment errors for most methods were concentrated near cortical-layer boundaries, where gradual transcriptional transitions make precise correspondence more difficult (Fig. 3c). PASTE exhibited additional mismatches within cortical layers. DeST-OT improved upon PASTE but still left many spots unmatched, whereas SLAT and Spateo aligned one slice reasonably well but left a substantial number of spots in the other slice unmatched. OT-knn produced fewer mismatched and unmatched spots across both slices than the comparison methods.

To further assess alignment quality, we used one slice in each pair to infer domain labels for the other slice using the best-matching pairs (Fig. 3d). Accurate alignment should yield inferred labels that closely resemble the original annotations. OT-knn produced inferred domain labels that closely resembled the original annotations in both directions. In contrast, PASTE showed substantial deviations from the annotated layers, and DeST-OT produced noticeable misassignments, particularly among layers 4–6, which have relatively similar gene-expression profiles [5]. SLAT and Spateo yields reasonable label transfer in one direction but substantially poorer results in the reverse direction.

The correspondence visualization for the A–B pair of Sample I provided a complementary view of spatial consistency (Fig. 3e). OT-knn produced coherent correspondences between anatomically similar regions, with relatively few crossing connections. DeST-OT showed more oblique and crossing correspondences and left a substantial number of spots unmatched. Spateo produced relatively coherent matches for one slice but left many spots in the other slice unmatched, indicating asymmetric alignment. PASTE and SLAT exhibited the most pronounced crossing correspondences, consistent with lower spatial coherence; SLAT also left many spots unmatched.

#### Sensitivity to neighborhood size

Finally, we evaluated the sensitivity of pairwise OT-knn alignment to the neighborhood size *k* using the B–C pair of Sample III (Fig. 3f). Performance remained stable across a broad range of neighborhood sizes, from 5 to 360 neighbors, indicating that OT-knn is relatively insensitive to this parameter.

#### Multi-slice reconstruction and domain identification

Having established the accuracy and robustness of pairwise alignment, we next examined whether the resulting correspondences could support analyses across multiple tissue sections. We sequentially registered the four slices of DLPFC Sample III using correspondences between consecutive slice pairs and stacked the registered slices in anatomical order. The resulting three-dimensional reconstruction preserved cortical-layer organization and showed spatially coherent alignment across slices (Fig. 3g).

We also linked best-matching correspondences across consecutive slices to construct spot chains for multi-slice domain identification. Clustering the chain-level representations produced spatial domains that were consistent across slices and closely matched the annotated cortical layers (S1 Fig). Additional quantitative evaluation is provided in the Supplementary Materials. These results demonstrate that OT-knn-derived correspondences can support both three-dimensional reconstruction and domain identification across multiple tissue sections.

### Robust alignment of mouse brain aging data under donor heterogeneity (MERFISH)

To evaluate the applicability of OT-knn to imaging-based spatial transcriptomics and its robustness to biological heterogeneity, we applied it to a mouse brain aging dataset [7] generated using MERFISH [23]. The dataset contains 31 brain tissue samples from 12 donors spanning three age groups (4, 24, and 90 weeks), providing a challenging setting characterized by high spatial resolution, cellular heterogeneity, and inter-donor variation. We evaluated alignment in two settings: (i) between slices from the same donor and (ii) between slices from different donors sampled from similar anatomical regions.

Across both settings, OT-knn achieved the highest mapping probability accuracy and best-matching accuracy among PASTE, DeST-OT, SLAT, Spateo, and OT-knn (S3 Fig and Fig. 4a). As expected, alignment accuracy decreased for all methods in the cross-donor setting, reflecting the greater biological variability between individuals. Although OT-knn also showed a modest decrease, it remained the top-performing method, indicating greater robustness to inter-donor variation than the comparison methods.

**Fig 4.**
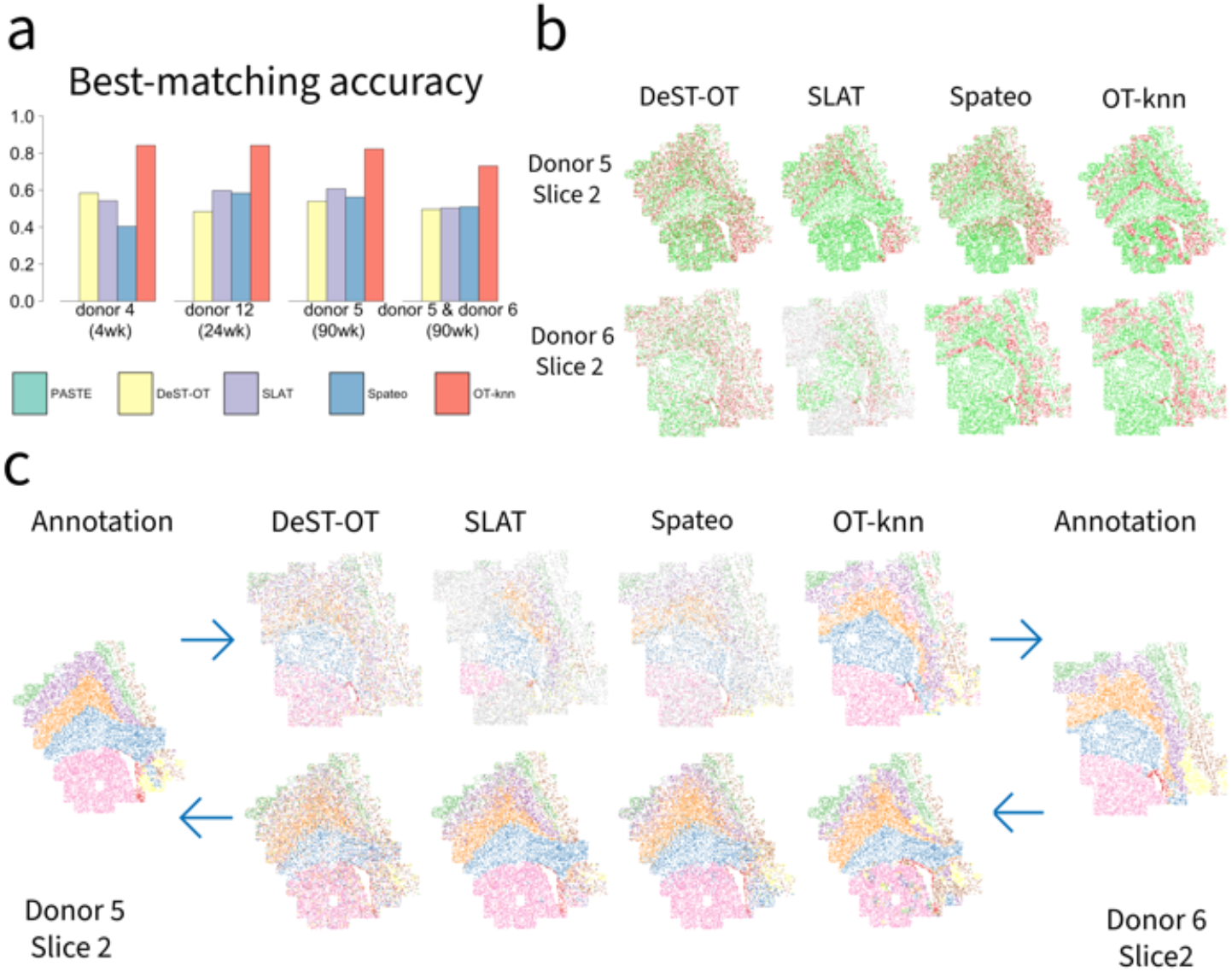
Performance comparison for mouse brain aging dataset. **a**. Best-matching accuracy for alignments of within-donor slice pairs from Donor 4 (4 weeks), Donor 12 (24 weeks), and Donor 5 (90 weeks), and a cross-donor slice pair between Donors 5 and 6 (90 weeks), comparing PASTE, DeST-OT, SLAT, Spateo, and OT-knn. **b**. Best-matching results for the cross-donor pair. Correctly aligned spots are shown in green, misaligned spots in red, and unaligned spots in gray. **c**. Domain inference results for the cross-donor pair by DeST-OT, SLAT, Spateo, and OT-knn.

Qualitative examination of the best-matching results further revealed differences in alignment behavior (Fig. 4b). Most mismatches produced by OT-knn were concentrated near spatial-domain boundaries. DeST-OT more frequently matched spots across distinct domains, whereas SLAT left many spots unmatched in one slice. Spateo produced alignment quality comparable to OT-knn in one direction but substantially poorer matching in the reverse direction, indicating asymmetric performance. Similar patterns were observed in bidirectional domain-label transfer (Fig. 4c): OT-knn preserved spatial-domain structure in both directions, whereas the comparison methods showed asymmetric or less accurate label inference. Overall, these results demonstrate that OT-knn provides accurate and robust alignment for imaging-based spatial transcriptomics data, including comparisons across donors.

### Alignment across developmental stages in the axolotl telencephalon (Stereo-seq)

We next evaluated whether OT-knn could align spatial transcriptomics (ST) slices across developmental stages, a challenging setting characterized by dynamic gene expression, tissue growth, and changes in cell-type composition. We analyzed an axolotl telencephalon dataset [17] generated using Stereo-seq [15]. The dataset spans five developmental stages, including three embryonic stages (Stages 44, 54, and 57), juvenile, and adult (Fig. 5a,b).

**Fig 5.**
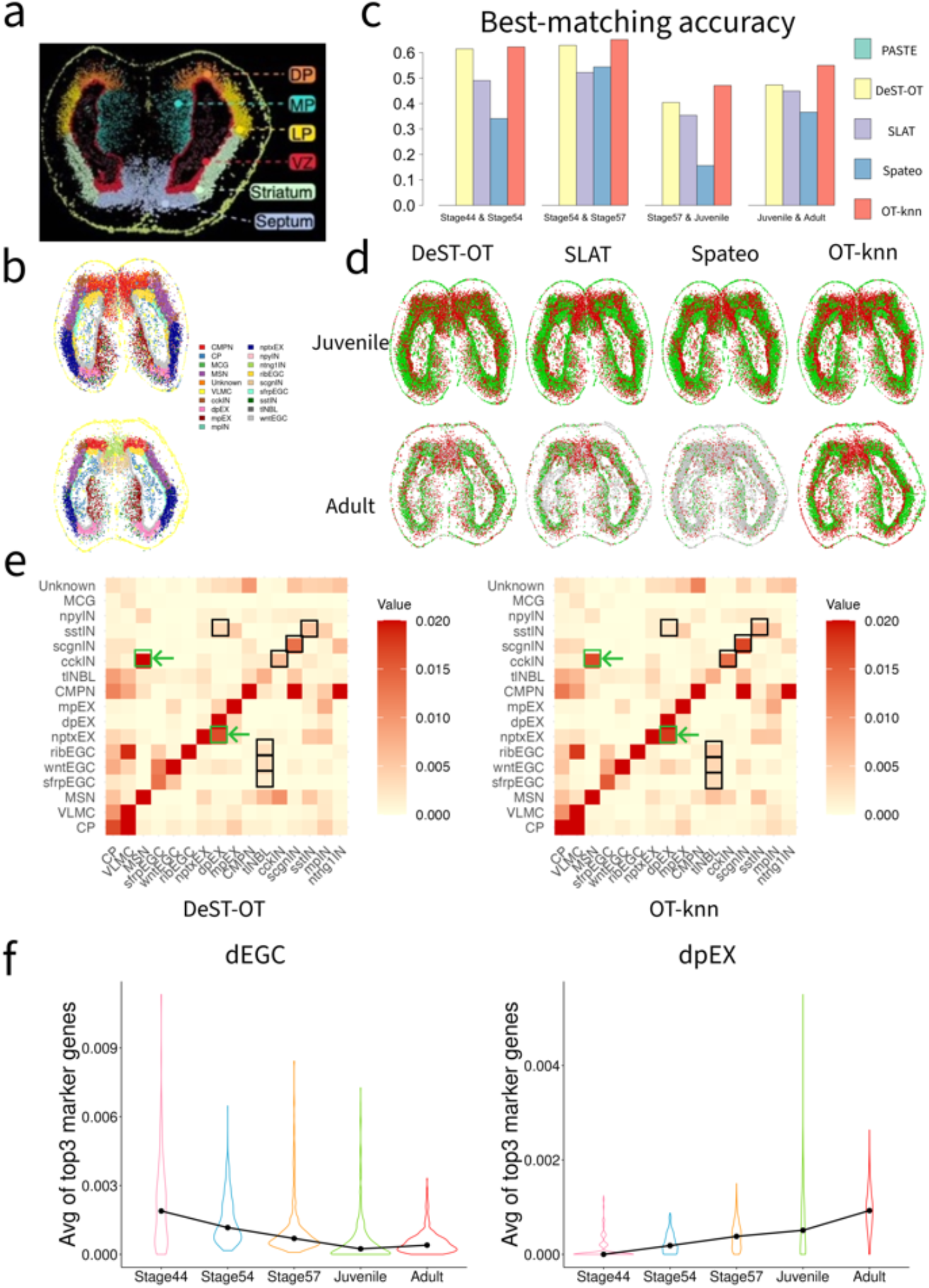
Analysis of axolotl telencephalon Stereo-seq data. **a**. Anatomical organization of the adult axolotl telencephalon identified by Stereo-seq [17]. DP, dorsal pallium; MP, medial pallium; LP, lateral pallium; VZ, ventricular zone. **b**. Cell-type annotations for the juvenile and adult axolotl telencephalon slices as defined in [17]. **c**. Best-matching accuracy for alignment between consecutive developmental stages using DeST-OT, SLAT, Spateo, and OT-knn. PASTE is excluded because it does not produce one-to-one spot matches. **d**. One-to-one alignment results between juvenile and adult slices. Correctly aligned spots are shown in green, misaligned spots in red, and unaligned spots in gray. **e**. Heatmap visualization of cell-type–level alignments between juvenile and adult slices produced by DeST-OT and OT-knn. Selected agreements and discrepancies referenced in the text are highlighted with green and black frames, respectively. **f**. Left: Average normalized marker gene expressions for dEGC across five developmental stages. Right: Average normalized marker gene expressions for dpEX across the same stages.

#### Cross-stage alignment performance

Because DeST-OT was specifically developed for aligning spatial transcriptomics data across developmental time points, it provides a particularly relevant benchmark in this setting. Across the cross-stage comparisons, OT-knn achieved the highest best-matching accuracy (Fig. 5c,d). It also achieved mapping probability accuracy comparable to or higher than that of DeST-OT, SLAT, and Spateo, while consistently outperforming PASTE (S6 Fig). These results indicate that OT-knn maintains reliable alignment despite substantial developmental changes in tissue structure and cellular composition.

#### Cell-type correspondence and developmental relationships

To examine alignment quality at the cell-type level, we calculated the average mapping probability between annotated cell types in the juvenile and adult slices and visualized the resulting values as heatmaps (Fig. 5e). Both OT-knn and DeST-OT exhibited a clear diagonal structure, indicating that corresponding cell types were generally preserved across stages. For several interneuron subtypes, however, OT-knn assigned higher within-cell-type mapping probabilities than DeST-OT. These included Cck^+^ inhibitory neurons (cckIN; 0.014 versus 0.006), Scgn^+^ inhibitory neurons (scgnIN; 0.017 versus 0.013), and Sst^+^ inhibitory neurons (sstIN; 0.007 versus 0.005), representing some of the largest differences observed along the diagonal.

The inferred mappings also captured biologically plausible relationships between related cell populations. Both OT-knn and DeST-OT aligned dpEX with nptxEX, two excitatory neuronal populations enriched in the pallium, and cckIN with MSN, two inhibitory neuronal populations enriched in the striatum. These correspondences are consistent with their shared neurotransmitter identities and spatial localization.

OT-knn more selectively emphasized several developmental relationships supported by prior biological knowledge. In particular, it assigned relatively high mapping probabilities between multiple ependymoglial cell (EGC) subtypes, including sfrpEGC, wntEGC, and ribEGC, and telencephalon neuroblasts (tlNBL). This pattern is consistent with the known transition from progenitor-like EGCs to neuroblasts in the axolotl telencephalon [17, 37]. DeST-OT assigned lower probabilities to these EGC–NBL relationships. Conversely, DeST-OT occasionally produced correspondences that were less consistent with known spatial or lineage relationships, such as alignment between dpEX and sstIN, which are spatially segregated and belong to distinct excitatory and inhibitory lineages [17]. Similar patterns were observed in the Stage 57–to–juvenile comparison, including alignment between immature MSN and nptxEX (S7 Fig).

#### Developmental marker-gene trajectories

To further assess the biological plausibility of the OT-knn-inferred developmental relationships, we sequentially applied OT-knn across the five developmental stages and examined marker-gene expression along the resulting cross-stage correspondences. We focused on development-associated ependymoglial cells (dEGCs) and mature excitatory neurons (dpEX). Previous studies have shown that EGCs function as neural progenitors that give rise to immature neurons, which subsequently differentiate into mature neurons during axolotl telencephalon development [17, 37].

As shown in Fig. 5f, expression of dEGC marker genes decreased progressively from embryonic to adult stages, whereas expression of dpEX marker genes increased over the same period. This opposing temporal pattern is consistent with the known progression from progenitor-like EGCs to mature excitatory neurons and supports the biological plausibility of the cross-stage correspondences inferred by OT-knn.

Overall, OT-knn preserved cell-type correspondence across developmental stages while also capturing biologically meaningful relationships among progenitor, immature, and mature neuronal populations. These findings demonstrate its robustness in settings involving substantial developmental changes in gene expression, tissue structure, and cellular composition.

### Cross-platform alignment between Stereo-seq and seqFISH mouse embryo datasets

To evaluate whether OT-knn can align spatial transcriptomics datasets generated by different technologies, we performed a cross-platform analysis using mouse embryo datasets profiled by Stereo-seq [15] and seqFISH [24, 36]. Cross-platform alignment is substantially more challenging than within-platform alignment because different technologies often vary in gene coverage, spatial resolution, measurement characteristics, and annotation granularity.

The Stereo-seq dataset corresponds to an E9.5 mouse embryo and contains approximately 5,000 cells with transcriptome-wide measurements covering more than 20,000 genes. The seqFISH dataset corresponds to an E8.75 mouse embryo and contains more than 10,000 cells but measures only 350 genes. In addition, the two datasets were annotated using different cell-type taxonomies. The seqFISH dataset contains 24 fine-grained cell types, whereas the Stereo-seq dataset contains 12 broader cell-type annotations. To facilitate quantitative evaluation, we mapped the original annotations from both datasets into seven shared major cell-type categories (Fig. 6a).

**Fig 6.**
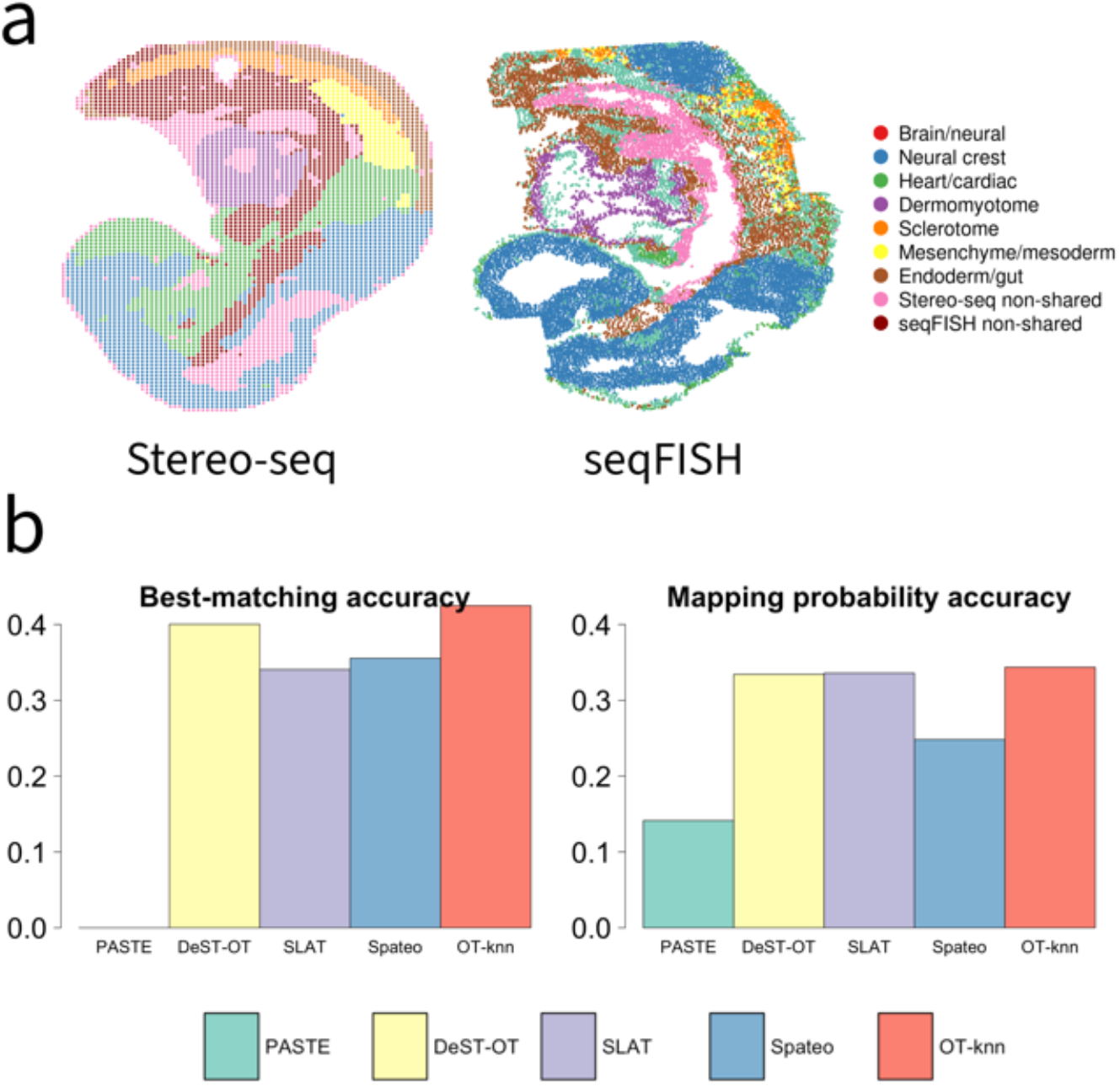
Performance comparison for mouse embryo datasets by Stereo-seq and seqFISH. **a**. Visualization of E9.5 (Stereo-seq) and E8.75 (seqFISH) mouse embryo datasets, colored by cell types. **b**. Best-matching accuracy and mapping probability accuracy for PASTE, DeST-OT, SLAT, Spateo, and OT-knn.

We compared OT-knn with PASTE, DeST-OT, Spateo, and SLAT using the same best-matching accuracy and mapping probability accuracy metric employed throughout the manuscript. Despite the substantial differences between the two platforms, OT-knn achieved the highest alignment accuracy among all methods evaluated (Fig. 6b). The best-matching accuracy of OT-knn was 0.42, exceeding those obtained by PASTE, DeST-OT, Spateo, and SLAT. Although the overall accuracy was lower than that observed in our within-platform analyses, this reduction is expected given the combined challenges of cross-platform measurement differences, distinct developmental stages, unequal gene coverage, and differences in annotation resolution.

These results demonstrate that OT-knn remains effective in the presence of substantial technological differences and suggest that neighborhood-aware representations can improve the robustness of spatial transcriptomics alignment beyond the same-platform setting.

### Computational efficiency

We assessed the computational efficiency of OT-knn across datasets spanning a wide range of sizes, with results summarized in Supplementary S1 Table. OT-knn completed all analyses on CPUs, with runtime increasing with the number of spots, and remained computationally practical for all datasets. Its runtime was broadly comparable to that of DeST-OT and was lower for several of the larger MERFISH comparisons. Among the other methods, Spateo generally required less CPU time, while SLAT was faster but relied on GPU acceleration. PASTE did not converge for most of the larger datasets under the evaluated settings. Overall, OT-knn showed practical computational performance across the datasets examined.

## Discussion

We introduce OT-knn, a neighborhood-aware method for aligning spatial transcriptomics (ST) slices by integrating local spatial context within an optimal transport framework. Rather than relying solely on individual spot-level expression profiles, OT-knn reconstructs each spot using information from its spatial neighbors, providing a more stable representation of the local tissue microenvironment before alignment.

Taken together, the simulation and real-data analyses demonstrate that OT-knn is robust to several major challenges in ST alignment, including geometric deformation, transcriptional noise, inter-individual heterogeneity, and developmental change. Across sequencing- and imaging-based technologies, the inferred correspondences preserved anatomical organization and biologically plausible cell-type relationships. OT-knn also remained effective across platforms, and its pairwise correspondences could be linked across serial sections to support three-dimensional reconstruction and multi-slice domain identification. This consistent performance likely arises from the incorporation of local spatial context: reconstructing each spot from neighboring expression profiles reduces sensitivity to sparse or noisy measurements and captures local tissue organization that may be missed by single-spot representations. Optimal transport then provides flexible probabilistic correspondences across slices. Together, these components support robust alignment under diverse sources of technical and biological variation.

One limitation of the current framework is its use of balanced OT, which requires the probability mass in each slice to be fully matched to the other. This assumption may lead to forced correspondences when tissue sections contain substantial non-overlapping regions. Extending OT-knn using unbalanced or partial optimal transport, which allows some probability mass to remain unmatched, is therefore an important direction for future work. A systematic treatment of partial overlap will require further investigation of overlap-aware modeling, parameter selection, and validation across diverse tissue geometries.

Scalability will become increasingly important as technologies such as Visium HD and Xenium generate datasets at near-single-cell or subcellular resolution. The sparse spatial *k*-nearest-neighbor graph in OT-knn can be constructed efficiently using approximate nearest-neighbor search [38]. As in other OT-based alignment methods, the main computational cost arises from the dense pairwise cost and transport matrices, which require *O*(*nm*) memory for slices containing *n* and *m* spots. Entropic regularization with Sinkhorn optimization [39] and low-rank OT approaches such as FRLC [40] offer practical directions for improving the computational and memory efficiency of this step.

Overall, OT-knn provides a flexible framework for aligning ST data across tissue sections, individuals, developmental stages, and measurement technologies, while supporting downstream cross-sample integration and three-dimensional tissue analysis.

## Methods

### Overview of the OT-knn framework

OT-knn aligns spatial transcriptomics slices by integrating local neighborhood information with optimal transport–based probabilistic matching. Given a pair of spatial transcriptomics slices, the method first constructs a neighborhood-aware representation of each spot and then performs probabilistic alignment across slices. Specifically, OT-knn consists of two main steps. First, each spot is embedded in its local tissue context by reconstructing its gene expression profile from a spatial *k*-nearest-neighbor graph, producing a microenvironment-aware representation that reduces sensitivity to noise and local variability. Second, optimal transport is applied to align the reconstructed profiles across slices, yielding a probabilistic mapping between spots that preserves global structure while allowing flexible correspondence. One-to-one matches can be extracted from this mapping when required. For analyses involving more than two slices, OT-knn can be applied sequentially to adjacent slice pairs, and the resulting pairwise correspondences can be linked across slices to support three-dimensional reconstruction and multi-slice domain identification. This framework enables robust pairwise and multi-slice analysis across datasets with spatial distortion, expression variability, and developmental differences.

### OT-knn algorithm

OT-knn takes a pair of spatial transcriptomics slices as input and first applies a unified preprocessing procedure to place the data in a shared, denoised low-dimensional expression space for alignment. Specifically, it (1) retains the genes shared between the two slices; (2) concatenates the count matrices and performs library-size normalization followed by log transformation; (3) conducts feature selection by choosing the top 5,000 highly variable genes when more than 5,000 genes are available, or all genes otherwise; and (4) applies principal component analysis (PCA), retaining the top 50 components as the expression representation for alignment.

After preprocessing, each slice is represented as a pair (*X, Z*), where *X* ∈ ℝ^*n*×*p*^ denotes the low-dimensional expression matrix with *n* spots and *p* principal components, and *Z* ∈ ℝ^*n*×2^ contains the spatial coordinates of the spots. Given two slices, OT-knn performs alignment using (*X*_1_, *Z*_1_) and (*X*_2_, *Z*_2_) as input.

#### Spatial neighbor graph and reconstruction of gene expression profiles

To incorporate local spatial context, we characterize the cellular microenvironment of each spot using its *k*-nearest neighbors (kNN). For a spot *i*, let *N*_*k*_(*i*) denote its *k* nearest neighbors under Euclidean distance in the spatial plane. Each neighbor *j* ∈ *N*_*k*_(*i*) is assigned a nonnegative weLight *w*_*ij*_ that decreases with spatial distance. The weights are normalized as 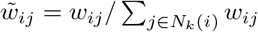.

We define the distance-based weighting scheme as:

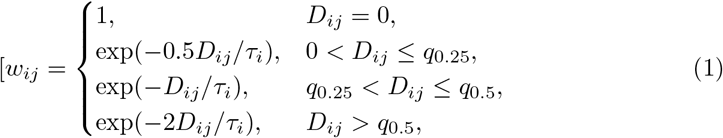

where *D*_*ij*_ is the Euclidean distance between spots *i* and *j*, and *ω*_*i*_ is the maximum distance between spot *i* and its *k* nearest neighbors. The thresholds *q*_0.25_ and *q*_0.5_ are defined as the 25th percentile and median of all kNN distances pooled across spots.

The reconstructed microenvironment expression profile for spot *i* is given by

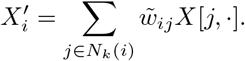

We denote the reconstructed expression matrices for the two slices as 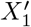 and 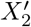.

For the analyses in this study, we set *k* = 100 for the DLPFC and mouse brain aging datasets and *k* = 20 for the human intestine, cross-platform mouse embryo, and axolotl telencephalon datasets. These choices reflect differences in spatial resolution and tissue heterogeneity across datasets. In general, we recommend setting *k* to approximately 1–5% of the total number of spots, with adjustment based on spatial resolution and tissue complexity.

#### Pairwise alignment via optimal transport

We use optimal transport (OT) to perform pairwise alignment between slices. OT provides a principled framework for matching two distributions by minimizing the total cost of transporting mass between them. In the context of ST alignment, OT allows each spot to distribute its matching probability across multiple candidate spots, naturally accommodating biological variability, noise, and differences in slice size, without enforcing a hard one-to-one correspondence.

To align the reconstructed expression profiles 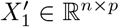 and 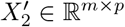, we solve the Earth Mover’s Distance (EMD) problem [41]:

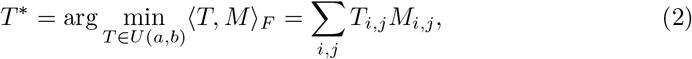

subject to

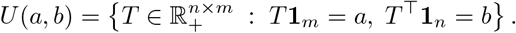

Here,

- *M* ∈ ℝ^*n* ×*m*^ is the gene expression dissimilarity matrix, where 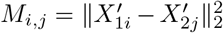 is the squared Euclidean distance between reconstructed profiles.
- 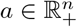 and 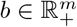 are the marginal distributions over spots in the two slices, satisfying 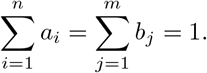

In this study, we use uniform marginals, i.e., *a*_*i*_ = 1*/n* and *b*_*j*_ = 1*/m*, assigning equal weight to all spots.
- *T* ^∗^ is the optimal transport plan, where 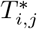 represents the probability mass transported from spot *i* in slice 1 to spot *j* in slice 2.

### Evaluation metrics

We treated the annotation of domain labels provided in each dataset as the ground truth, and evaluated alignment performance using two metrics: mapping probability accuracy and best-matching accuracy.

#### Mapping probability accuracy

This metric, originally introduced in PASTE [29], quantifies the *overall* quality of probabilistic alignment by summing the mapping probabilities of spot pairs that belong to the same domain label. Formally, let *T* = [*T*_*i,j*_] denote the normalized pairwise mapping probability matrix between spots in two slices, and let *l*(*i*) denote the domain label of spot *i*. Then mapping probability accuracy is defined as:

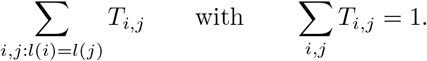

A higher value indicates that most of the mapping probability mass is assigned to spot pairs that share the same domain label. It captures the distribution of probabilistic alignment mass across all potential matches.

#### Best-matching accuracy

This metric evaluates the quality of one-to-one spot alignments. For each spot, we define its best match as the spot with the highest mapping probability:

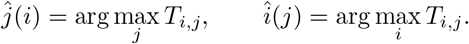

The best-matching accuracy is defined as the average proportion of correctly aligned spots across the two slices:

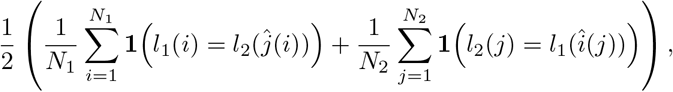

where *l*_1_(*i*) and *l*_2_(*j*) denote the domain labels of spots in slices 1 and 2, respectively, 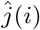 and 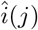 denote their matched counterparts, and **1**(·) is the indicator function. Here, *N*_1_ and *N*_2_ are the numbers of spots whose domain labels are shared between the two slices. Spots belonging to domains present in only one slice are excluded, as they do not admit meaningful cross-slice correspondences. This restriction ensures that the accuracy metric reflects alignment quality only for comparable spots.

Unlike mapping-probability accuracy, which evaluates the distribution of probabilistic alignment mass, best-matching accuracy focuses on one-to-one assignments. A higher best-matching accuracy therefore indicates a larger proportion of correctly aligned spots among those that have domain-consistent counterparts.

### Multi-slice reconstruction and domain identification

Consider an anatomically ordered set of *L* spatial transcriptomics slices, denoted by *S*_1_, …, *S*_*L*_. OT-knn is first applied to each pair of consecutive slices, (*S*_1_, *S*_2_), …, (*S*_*L*−1_, *S*_*L*_), to obtain pairwise probabilistic mappings and best-matching spot correspondences. For three-dimensional reconstruction, the correspondences between consecutive slices are used sequentially to register the slices, which are then stacked according to their anatomical order to construct a three-dimensional tissue representation.

For multi-slice domain identification, the best-matching correspondences from consecutive slice pairs are linked transitively to form spot chains that trace corresponding locations across the ordered slices. After restricting the data to genes shared across all slices and applying the preprocessing described above, the expression matrices from all slices are concatenated by rows. Principal component analysis (PCA) is then performed on the concatenated expression matrix, yielding a common latent representation for all spots. For each spot chain, the PCA embeddings of its constituent spots are averaged to obtain a chain-level representation. *K*-means clustering is applied to the chain-level representations, and the resulting cluster label for each chain is assigned back to all corresponding spots across slices.

### Simulation design

To generate simulated ST data with known ground-truth alignment, we used a real ST slice as the reference and constructed a second, perturbed slice under four simulation scenarios: (1) nonlinear spatial distortion, (2) gene expression variation, and (3) a combined perturbation incorporating both (1) and (2). To assess robustness under a more challenging and realistic setting, we additionally considered (4) a composite perturbation combining global slice misalignment, local non-rigid deformation, gene expression variation, and spot dropout.

Specifically, we used an adult human intestine tissue slice profiled with the 10x Genomics Visium platform [21, 35], which contains 2,649 spots and expression measurements for 33,538 genes, as the reference slice. In scenario (1), nonlinear spatial distortion was introduced by applying a mild nonlinear transformation to the spatial coordinates, (*x*^1.02^, *y*^1.01^). In scenario (2), gene expression variation was simulated by scaling gene counts at each spot by a random factor between 0.7 × and 1.3 ×, followed by the addition of small random noise (0–10 counts). Scenario (3) applied both perturbations simultaneously.

Scenario (4) introduced multiple spatial and transcriptional perturbations simultaneously. To mimic global misalignment between tissue sections, we first applied a 45^◦^ rotation together with translations corresponding to 20% and 15% of the tissue width and height, respectively. To model local non-rigid deformation, we divided the tissue into a 5 × 5 grid and applied independent random rotations of up to 6^◦^ and translations of up to 2% of the tissue size to each patch. We then introduced gene expression variation using the same scaling and noise model as in scenario (2) and preferentially removed low-count spots to mimic missing or low-quality capture locations. Because all perturbed slices were generated directly from the reference slice, the true spot-to-spot correspondence was known for every spot in scenarios (1)–(3) and for all retained spots in scenario (4).

### Benchmarking methods

We benchmarked OT-knn against PASTE, DeST-OT, Spateo, and SLAT across all datasets using the default parameter settings recommended in the original publications. Method-specific preprocessing was applied only when required. For PASTE, we retained genes with at least 100 total counts per slice for the DLPFC dataset and 15 for the axolotl dataset, following the authors’ guidelines. For all datasets, we ran PASTE using its standard workflow and default initialization of the transport plan. This differs from the publicly available implementation used for the PASTE analysis of the DLPFC dataset, which employs a dataset-specific, geometry-informed initialization constructed from the spatial coordinates of the paired slices. This tailored initialization can be advantageous when the slices have similar global geometry, orientation, and anatomical coverage, but it may be less broadly applicable when these properties differ. Moreover, the other benchmarked methods do not use comparable dataset-specific initialization. We therefore used the standard PASTE workflow to maintain a consistent and non-tailored benchmarking procedure across methods and datasets. For Spateo, we retained genes with at least 3 total counts per slice, following its recommended preprocessing. For OT-knn, genes with low expression (total counts *<* 15 within a slice) were removed for the DLPFC and axolotl datasets. DeST-OT and SLAT were run on the unfiltered data, as these methods are designed to operate directly on raw inputs. To ensure comparability across methods, we did not remove low-count spots for any method. Because SLAT is order-dependent, we consistently placed the slice with fewer spots as the source slice.

PASTE, DeST-OT, and Spateo produce a probabilistic mapping matrix *T* = [*T*_*ij*_] that quantifies the alignment probability between spots across slices. For these methods, one-to-one correspondences were obtained by assigning each spot to the counterpart with the highest mapping probability. SLAT directly outputs spot embeddings and one-to-one matching pairs but does not provide a probabilistic mapping matrix. Therefore, all best-matching accuracy evaluations and matching visualizations for SLAT were based on its original matching output. To enable evaluation using mapping probability accuracy, which requires a probabilistic mapping matrix, we additionally constructed a probability matrix from the SLAT embeddings by computing an optimal transport plan between the two embedding spaces, minimizing the Wasserstein distance as in Equation 2. This OT-derived probability matrix was used only for calculating mapping probability accuracy and did not affect SLAT’s matching results and visualizations.

### Marker gene trajectory analysis across developmental stages for axolotl brain dataset

To examine the biological relevance of the developmental trajectories inferred from alignment, we analyzed marker gene expression dynamics for development-related ependymoglial cells (dEGCs) and mature excitatory neurons (dpEX), which are expected to exhibit progenitor-to-neuron transitions during axolotl telencephalon development [17, 37]. As the original study [17] identified TMIE, ADCYAP1, and TIAM1 as the top three marker genes for dpEX and FABP7[nr], SLC1A3, and GFAP as the top three marker genes for dEGC, we tracked their expression levels across the developmental stages. For each spatial spot, marker gene expression was normalized by computing the percentage of each marker gene relative to the total expression within that spot. For each cell type, the normalized values of its three marker genes were averaged to obtain a single summary expression score per spot.

To track dpEX trajectories across development, we initiated the analysis at the adult stage by selecting spots annotated as dpEX in the original study [17]. Corresponding spots in earlier stages were identified through OT-knn alignments, enabling reconstruction of matched spot trajectories spanning Stage 44 through the adult stage. The distributions of average normalized marker expression for dpEX and dEGC were visualized across developmental stages using violin plots. To summarize temporal trends, the median value at each stage was computed and connected across stages.

## Supporting information

Supplementary Materials: OT-knn: a neighborhood-aware optimal transport framework for aligning spatial transcriptomics data

## Data availability

The human intestine dataset is available using GEO accession number GSE158328. The DLPFC dataset is available in the spatialLIBD package (https://research.libd.org/spatialLIBD/). The adult mouse brain dataset is available using GEO accession number GSE147747. The MERFISH dataset is available at https://cellxgene.cziscience.com/collections/31937775-0602-4e52-a799-b6acdd2bac2e. The Axolotl brain dataset is available at https://db.cngb.org/stomics/artista/. The cross-platform mouse embryo datasets are available from the MOSTA database and the Spatial Mouse Atlas resource: https://db.cngb.org/stomics/mosta/ and https://marionilab.cruk.cam.ac.uk/SpatialMouseAtlas/.

## Code availability

OT-knn is publicly available as a Python package at https://github.com/qunhualilab/OT_knn.

## Supporting information

**S1 Fig. Domain-identification results for the four slices of DLPFC Sample III**.

**S2 Fig. Mapping probability accuracy for the three samples in the human DLPFC dataset, comparing PASTE, DeST-OT, SLAT, Spateo, and OT-knn**.

**S3 Fig. Performance comparison for mouse brain aging dataset. a**. Mapping probability accuracy for slices from Donor 4 (4wk), Donor 12 (24wk), Donor 5 (90wk), and one pair of different donors, comparing PASTE, DeST-OT, SLAT, Spateo, and OT-knn. **b-d**. Best-matching results for the two slices of Donor 4, 5 and 12. Correctly aligned spots are shown in green, misaligned spots in red, and unaligned spots in gray.

**S4 Fig. Domain inference results for Donor 4, 5 and 12 in mouse brain aging dataset by DeST-OT, SLAT, and OT-knn**.

**S5 Fig. Visualization of the alignment for mouse brain aging dataset. Aligned spots are colored according to their annotated layers and unaligned spots are shown in gray**.

**S6 Fig. Mapping probability accuracy for alignment between consecutive time points of axolotl brain dataset using PASTE, DeST-OT, SLAT, Spateo, and OT-knn**.

**S7 Fig. Heatmap visualization of the alignment between slices of a. Stage 44 & Stage 54, b. Stage 54 & Stage 57 and c. Stage 54 & juvenile of axolotl brain dataset using DeST-OT and OT-knn**.

**S8 Fig. Performance comparison for adult mouse brain dataset. a**. Best-matching accuracy and mapping probability accuacy of five slice pairs for PASTE, DeST-OT, SLAT, and OT-knn. **b**. Mapping probability accuracy and best-matching accuracy across different numbers of nearest neighbors for one slice pair of Mouse A1. **c**. Best-matching results for three pairs from mouse A1 and A2. Correctly aligned spots are shown in green, misaligned spots in red, and unaligned spots in gray.

**S1 Table. Computational efficiancy**. Running time in seconds for the DLPFC (10x Genomics Visium), mouse brain aging (MERFISH), axolotl telencephalon (Stereo-seq), and adult mouse brain (Illumina NextSeq 500) datasets using PASTE, DeST-OT, SLAT, Spateo, and OT-knn. PASTE, DeST-OT, Spateo, and OT-knn were executed on CPUs, whereas SLAT was executed using GPU acceleration. “NA” indicates that the method did not converge within the specified maximum number of iterations.

