## Supplementary Materials: OT-knn: a neighborhood-aware optimal transport framework for aligning spatial transcriptomics data for "OT-knn: a neighborhood-aware optimal transport framework for aligning spatial transcriptomics data"

### Sensitivity to neighborhood size

We evaluated the sensitivity of OT-knn to the neighborhood size by varying $k$ from 5 to 360 using the B–C slice pair from Sample II of the DLPFC dataset. Both alignment metrics improved as $k$ increased from very small values and stabilized at approximately $k = 60$, remaining similar across a broad range of larger neighborhood sizes.

This stability is supported by the distance-weighted aggregation used in OT-knn, which assigns greater influence to nearby spots than to more distant neighbors. Consequently, even when $k$ is large, the reconstructed expression profile remains primarily determined by the local tissue context rather than approaching a global average. Overall, the performance is largely insensitive to further increases once $k$ is large enough, with no substantial performance decline attributable to over-smoothing over the range examined.

### Multi-slice analysis of DLPFC Sample III

We applied the multi-slice procedure described in the Methods to the four anatomically ordered slices (A–D) of DLPFC Sample III. OT-knn was first applied to the three consecutive slice pairs, A–B, B–C, and C–D, to obtain pairwise probabilistic mappings and best-matching spot correspondences.

For domain identification, best-matching correspondences from the three consecutive alignments were linked transitively to construct spot chains across the four slices. Only chains containing one spot from every slice were retained. Following the preprocessing described in the Methods, the expression matrices from the four slices were concatenated and projected into a common latent space using principal component analysis (PCA). The top 30 principal components were retained. For each spot chain, the PCA embeddings of its constituent spots were averaged to obtain a chain-level representation.

We examined clustering solutions with $K = 5 - 7$. The $K = 5$ solution produced more spatially coherent domains that were consistently represented across slices, whereas the $K = 6$ and $K = 7$ solutions included some clusters that spanned multiple annotated cortical layers. We therefore used $K = 5$ for the subsequent visualization and quantitative evaluation. This empirically selected value was not intended to impose a one-to-one correspondence between the inferred expression domains and the six annotated cortical layers plus white matter, because transcriptionally similar anatomical regions may be grouped into the same expression-derived domain.

The inferred domains were spatially coherent across the four slices and showed substantial agreement with the annotated cortical layers (Fig. S1). Quantitative comparison with the annotations yielded an adjusted Rand index (ARI) of 0.577 and a normalized mutual information (NMI) of 0.672. For context, Zhou et al. [1] reported an

ARI of 0.614 for STAligner, which was developed specifically for joint multi-slice integration, compared with 0.364 for PASTE on the same DLPFC sample. Because the analyses may differ in preprocessing, clustering, and parameter selection, these values are provided as contextual information rather than as a direct head-to-head comparison. Overall, these results indicate that correspondences obtained from consecutive OT-knn alignments preserve sufficient cross-slice structure to support domain identification across multiple tissue sections.

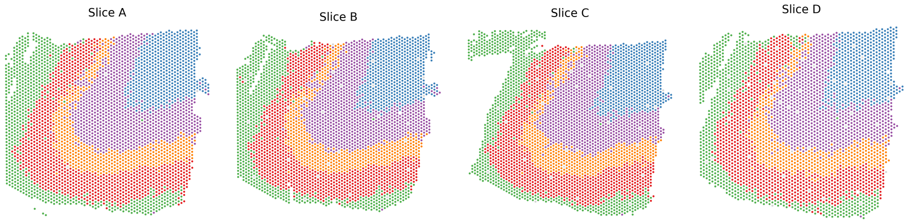


**S1 Fig. Domain-identification results for the four slices of DLPFC Sample III.**

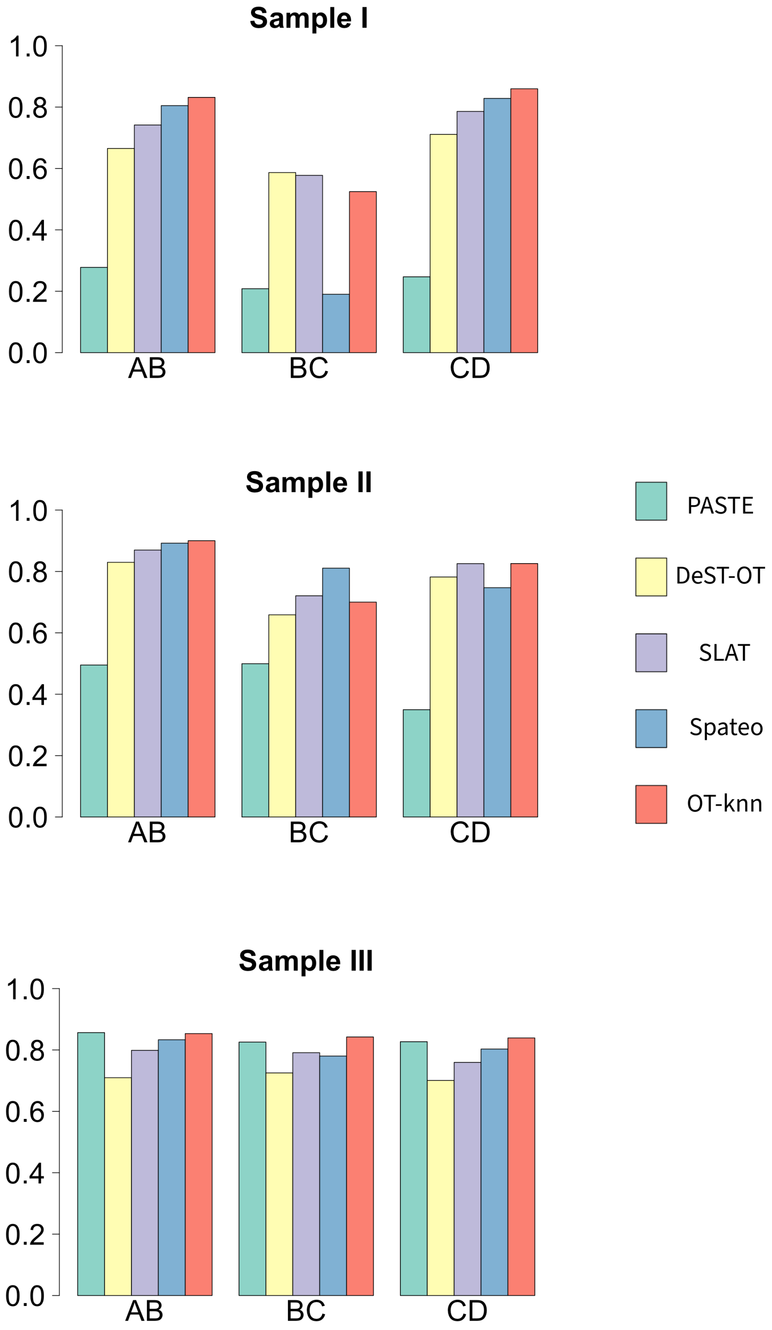


**S2 Fig. Mapping probability accuracy for the three samples in the human DLPFC dataset, comparing PASTE, DeST-OT, SLAT, Spateo, and OT-knn.**

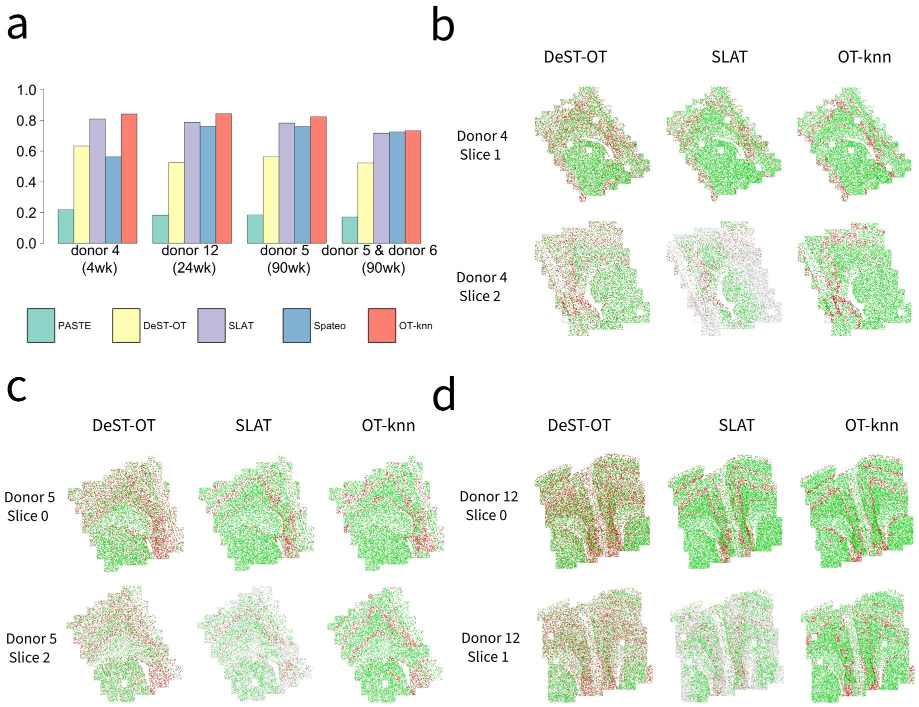


**S3 Fig. Performance comparison for mouse brain aging dataset. a.** Mapping probability accuracy for slices from Donor 4 (4wk), Donor 12 (24wk), Donor 5 (90wk), and one pair of different donors, comparing PASTE, DeST-OT, SLAT, Spateo, and OT-knn. **b-d**. Best-matching results for the two slices of Donor 4, 5 and 12. Correctly aligned spots are shown in green, misaligned spots in red, and unaligned spots in gray.

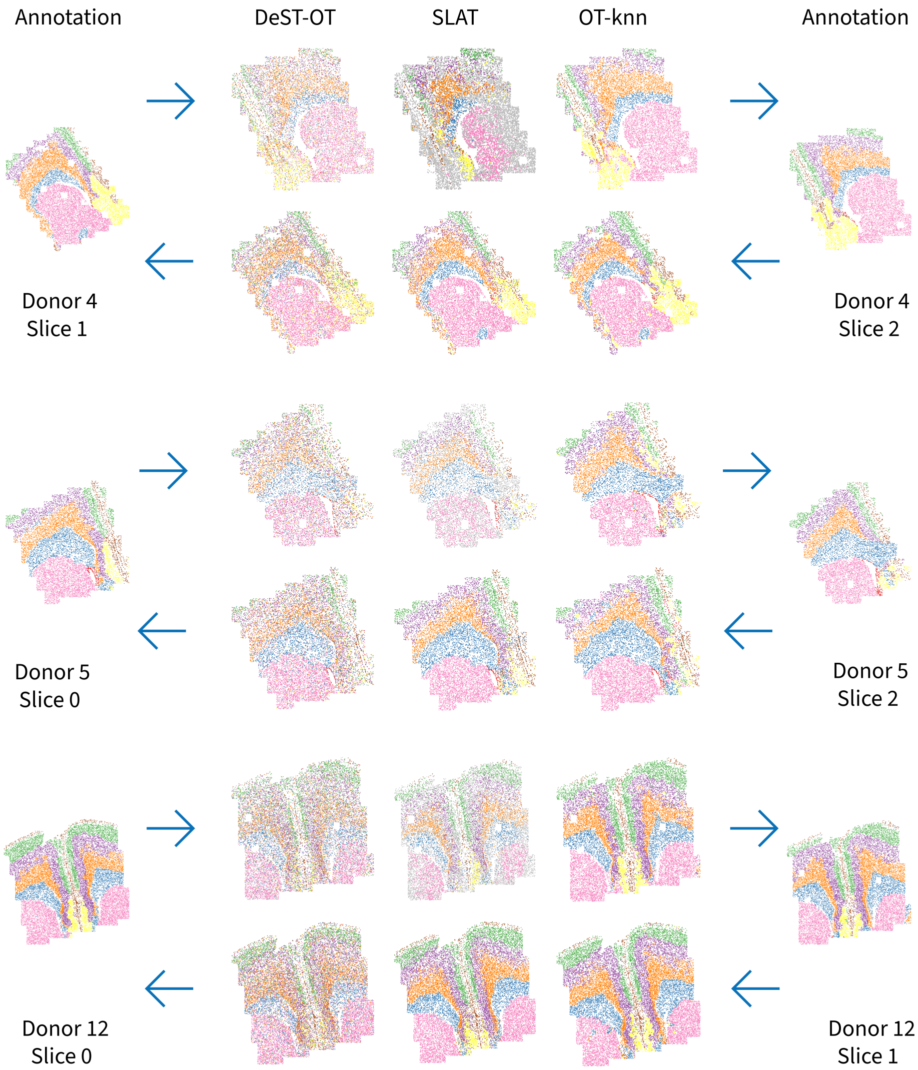


**S4 Fig. Domain inference results for Donor 4, 5 and 12 in mouse brain aging dataset by DeST-OT, SLAT, and OT-knn.**

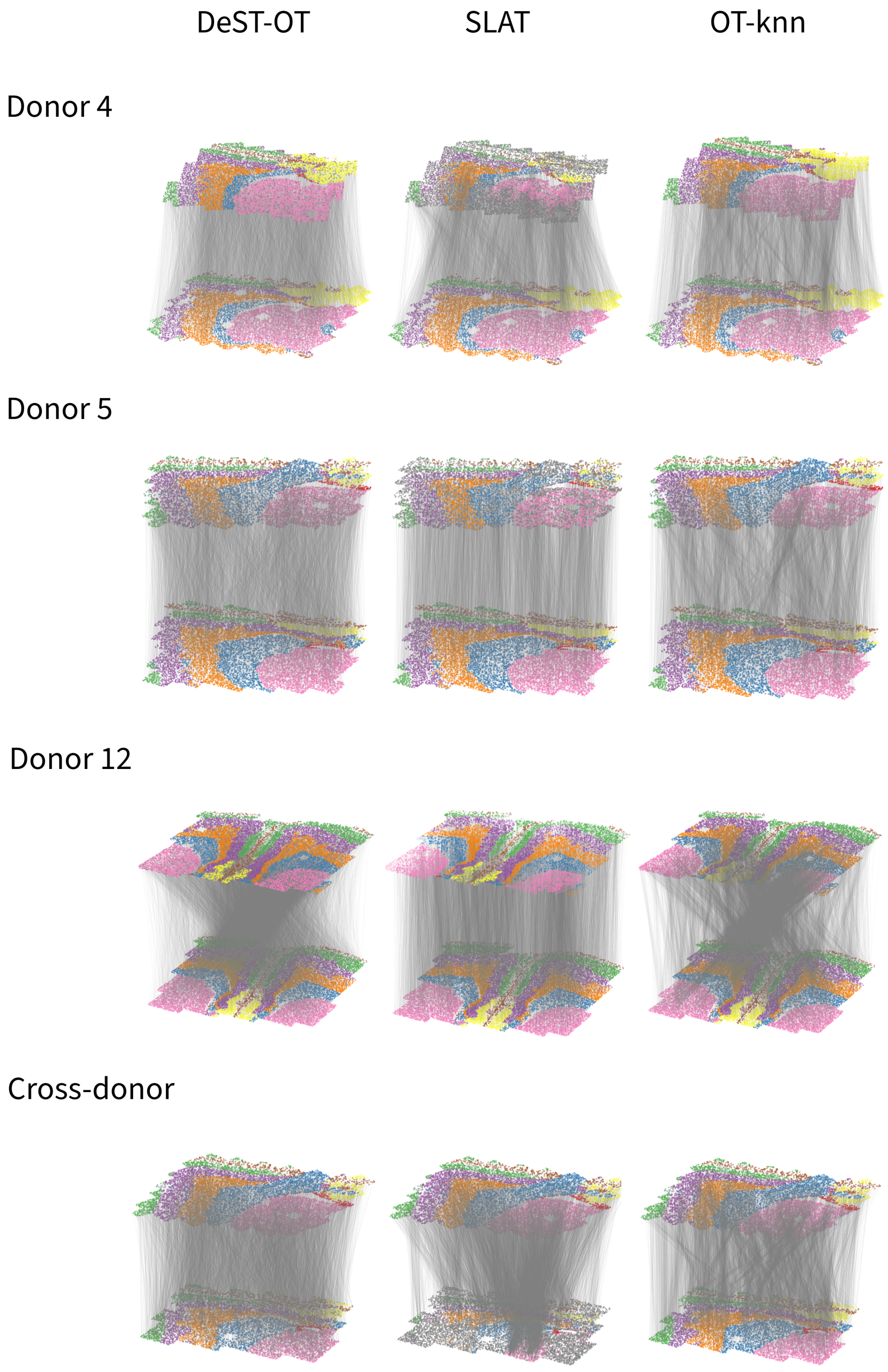


**S5 Fig. Visualization of the alignment for mouse brain aging dataset. Aligned spots are colored according to their annotated layers and unaligned spots are shown in gray.**

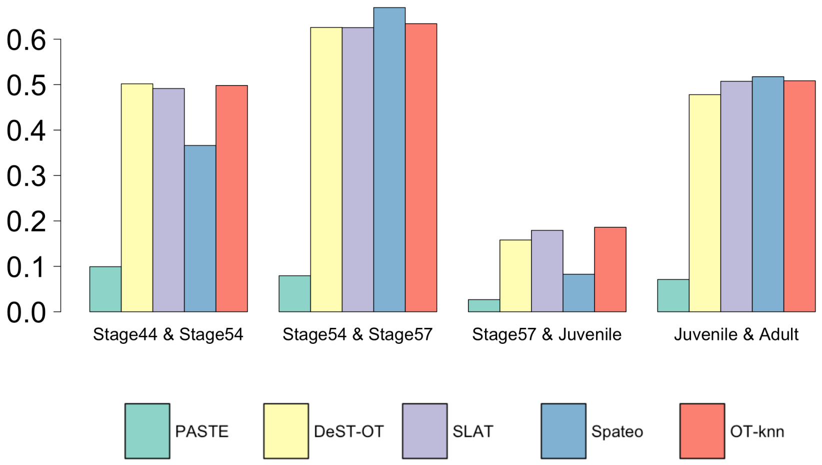


**S6 Fig. Mapping probability accuracy for alignment between consecutive time points of axolotl brain dataset using PASTE, DeST-OT, SLAT, Spateo, and OT-knn.**

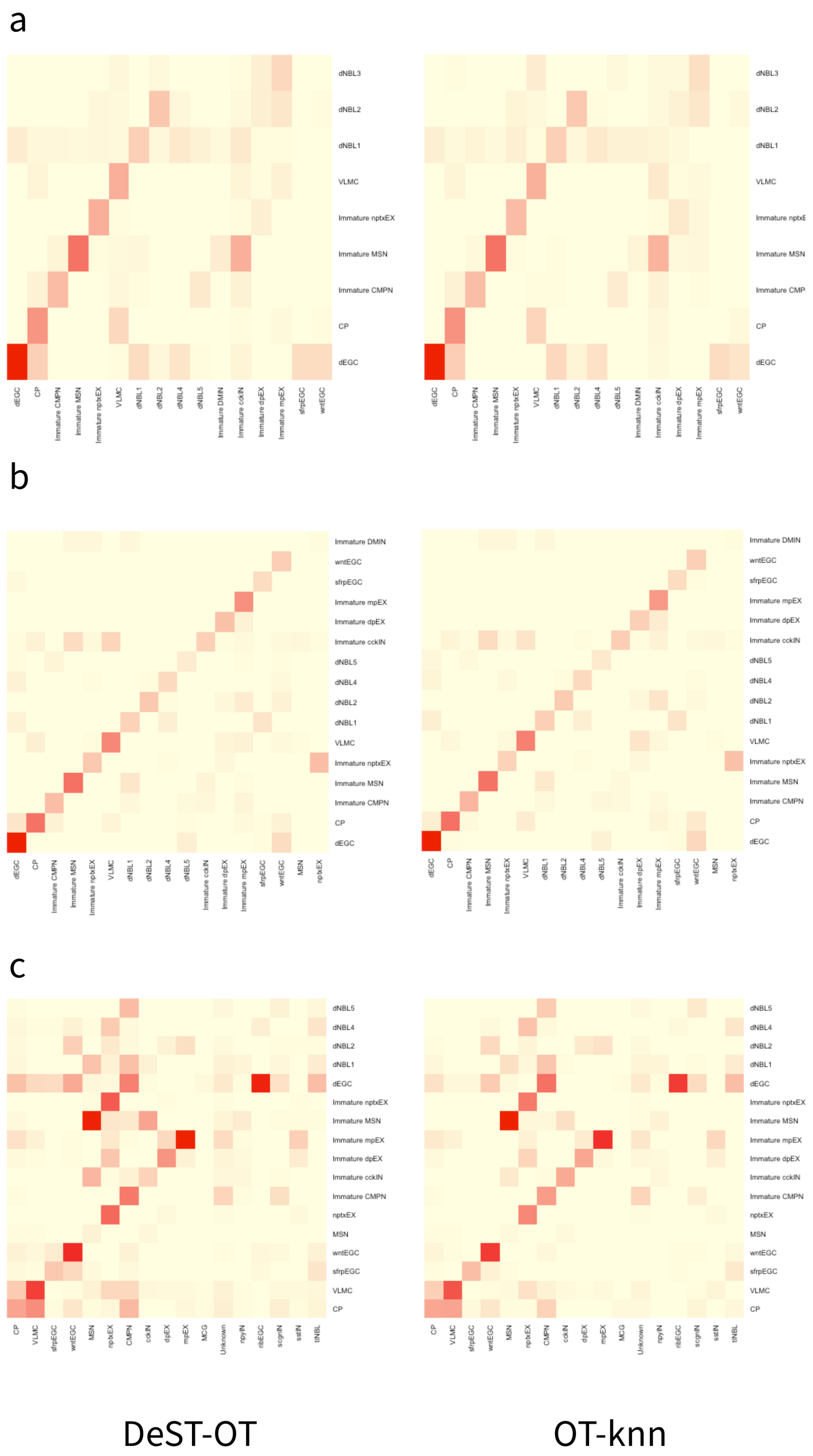


**S7 Fig. Heatmap visualization of the alignment between slices of a. Stage 44 & Stage 54, b. Stage 54 & Stage 57 and c. Stage 54 & juvenile of axolotl brain dataset using DeST-OT and OT-knn.**

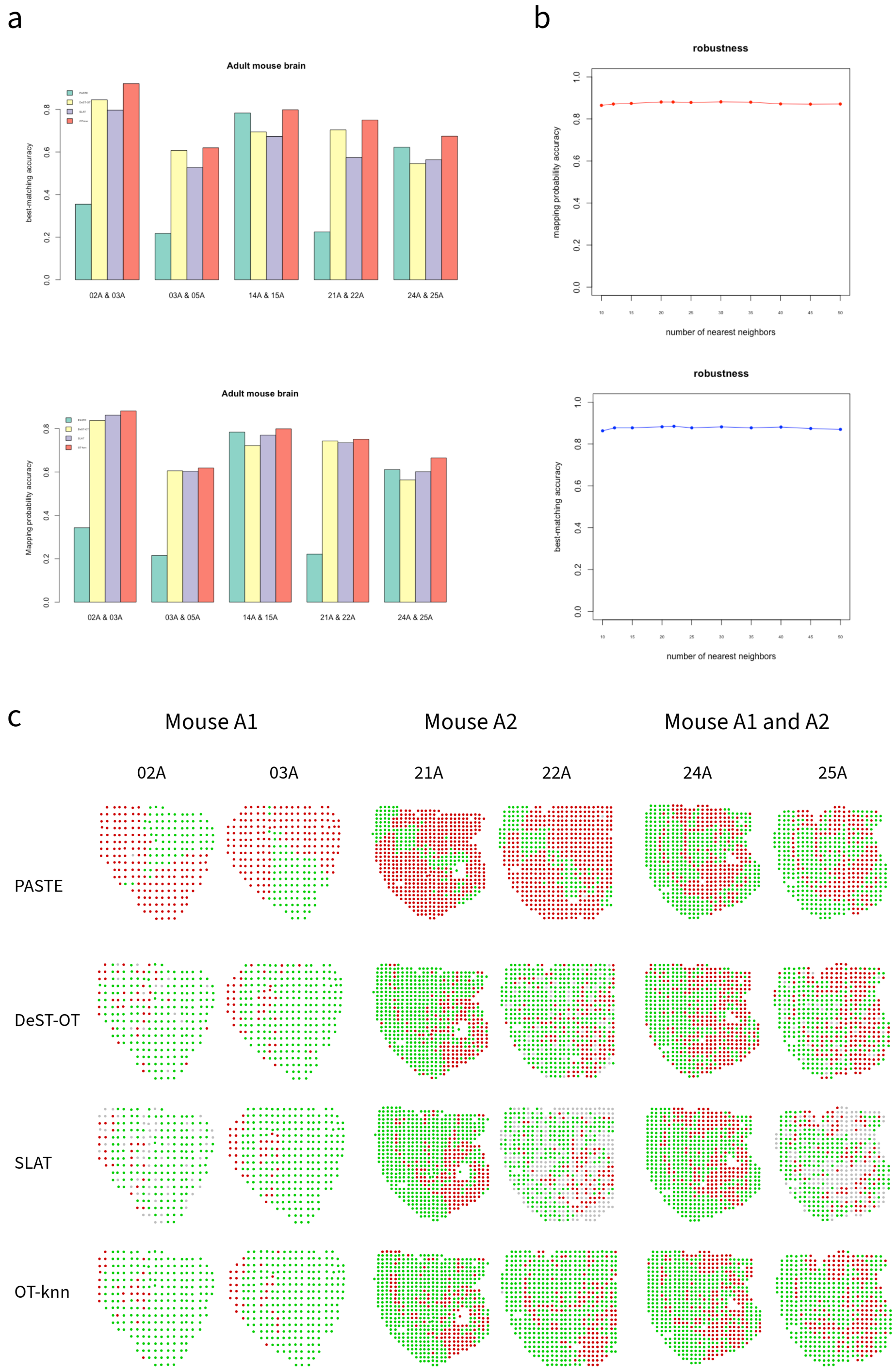


**S8 Fig. Performance comparison for adult mouse brain dataset. a.** Best-matching accuracy and mapping probability accuacy of five slice pairs for PASTE, DeST-OT, SLAT, and OT-knn. **b.** Mapping probability accuracy and best-matching accuracy across different numbers of nearest neighbors for one slice pair of Mouse A1. **c.** Best-matching results for three pairs from mouse A1 and A2. Correctly aligned spots are shown in green, misaligned spots in red, and unaligned spots in gray.

| Dataset | PASTE | DeST-OT | SLAT(GPU) | Spateo | OT-knn |
|---|---|---|---|---|---|
| **DLPFC** | | | | | |
| Spots (3k–4k) | | | | | |
| 151507 & 151508 | NA | 194 | 17 | 60 | 202 |
| 151508 & 151509 | NA | 229 | 18 | 68 | 212 |
| 151509 & 151510 | NA | 239 | 18 | 62 | 219 |
| 151669 & 151670 | NA | 124 | 14 | 45 | 126 |
| 151670 & 151671 | NA | 144 | 13 | 55 | 145 |
| 151671 & 151672 | NA | 157 | 14 | 53 | 167 |
| 151673 & 151674 | NA | 125 | 13 | 46 | 149 |
| 151674 & 151675 | NA | 124 | 12 | 46 | 129 |
| 151675 & 151676 | NA | 119 | 13 | 44 | 128 |
| **Mouse brain (MERFISH)** | | | | | |
| Spots (8k–17.5k) | | | | | |
| Donor 4 (4 wk) | NA | 2207 | 28 | 144 | 1261 |
| Donor 12 (24 wk) | NA | 7284 | 59 | 374 | 3036 |
| Donor 5 (90 wk) | NA | 1610 | 24 | 142 | 970 |
| Donor 5 & Donor 6 (90 wk) | NA | 1100 | 19 | 100 | 814 |
| **Axolotl brain** | | | | | |
| Spots (1.4k–11.7k) | | | | | |
| Stage 44 & 54 | NA | 41 | 6 | 31 | 61 |
| Stage 54 & 57 | NA | 119 | 10 | 61 | 144 |
| Stage 57 & Juvenile | NA | 788 | 20 | 159 | 776 |
| Juvenile & Adult | NA | 1677 | 27 | 149 | 1013 |
| **Adult mouse brain** | | | | | |
| Spots (240–618) | | | | | |
| 02A (A1) & 03A (A1) | 0.56 | 1.42 | 1.8 | 1.21 | 1.21 |
| 03A (A1) & 05A (A1) | 1.34 | 1.77 | 2.22 | 1.33 | 1.72 |
| 14A (A1) & 15A (A1) | 1.24 | 3.59 | 3.62 | 3.91 | 4.17 |
| 21A (A2) & 22A (A2) | 5.53 | 3.46 | 3.35 | 3.7 | 4.82 |
| 24A (A2) & 25A (A1) | 2.97 | 3.11 | 2.46 | 3.41 | 4.22 |

**S1 Table. Computational efficiency.** Running time in seconds for the DLPFC (10x Genomics Visium), mouse brain aging (MERFISH), axolotl telencephalon (Stereo-seq), and adult mouse brain (Illumina NextSeq 500) datasets using PASTE, DeST-OT, SLAT, Spateo, and OT-knn. PASTE, DeST-OT, Spateo, and OT-knn were executed on CPUs, whereas SLAT was executed using GPU acceleration. "NA" indicates that the method did not converge within the specified maximum number of iterations.
